# Precise gene models using long-read sequencing reveal a unique poly(A) signal in *Giardia lamblia*

**DOI:** 10.1101/2021.04.06.438666

**Authors:** Danielle Y. Bilodeau, Ryan M. Sheridan, Balu Balan, Aaron R. Jex, Olivia S. Rissland

## Abstract

During pre-mRNA processing, the poly(A) signal is recognized by a protein complex that ensures precise cleavage and polyadenylation of the nascent transcript. The location of this cleavage event establishes the length and sequence of the 3′ UTR of an mRNA, thus determining much of its post-transcriptional fate. Here, using long-read sequencing, we characterize the polyadenylation signal and related sequences surrounding *Giardia lamblia* cleavage sites for over 2600 genes. We find that *G. lamblia* uses a AGURAA poly(A) signal, which differs from the mammalian AAUAAA. We also describe how *G. lamblia* lacks common auxiliary elements found in other eukaryotes, along with the proteins that recognize them. Further, we identify 133 genes that show evidence of alternative polyadenylation. These results suggest that despite pared down cleavage and polyadenylation machinery, 3′ end formation still appears to be an important regulatory step for gene expression in *G. lamblia*.

## INTRODUCTION

Pre-mRNA processing is central to the proper expression and function of a gene. In eukaryotes, capping, splicing, and cleavage and polyadenylation of a pre-mRNA must all occur before export to the cytoplasm, and errors at any of these steps can have important consequences on post-transcriptional regulation. One key processing step is cleavage and polyadenylation. Here, the nascent RNA is cleaved at a precise location, which establishes the 3′ end of the mature transcript, and a poly(A) tail is added, which is required for downstream events in gene expression (Gallie, 1991; Singh et al., 2015). Adding to this complexity, some genes contain more than one cleavage site, resulting in isoforms with different 3′ UTRs and often different post-transcriptional fates (Mayr & Bartel, 2009; Sandberg et al., 2008; Tian et al., 2005). Alternative polyadenylation (APA) is widespread in many species, including *S. cerevisiae* and *S. pombe*, and more than half of human and mouse genes have multiple mRNA cleavage sites (Hoque et al., 2013; Liu et al., 2017; Moqtaderi et al., 2018; Tian et al., 2005). Inappropriate cleavage and polyadenylation can have severe, widespread consequences for gene expression, and is associated with cancer and lethality (Whitelaw & Proudfoot, 1986; Morris et al., 2012; Nourse et al., 2020), highlighting the central importance of this processing step.

Cleavage and polyadenylation is a complex, highly coordinated step that must be highly specific and sensitive. In humans, this process involves 20 core proteins and several cis-acting elements in the mRNA (Kumar et al., 2019). The main sequence element that directs cleavage is the polyadenylation signal (known as the poly(A) signal), which is an AAUAAA hexamer in metazoans (Proudfoot & Brownlee, 1976; Beaudoing, 2000). This hexamer and single-nucleotide variants, such as AUUAAA, are in turn recognized by a multi-protein complex known as the <u>c</u>leavage and <u>p</u>olyadenylation <u>s</u>pecificity <u>f</u>actor (or “CPSF”), which is composed of CPSF160, CPSF30, WDR33, CSPF73, CPSF100, Symplekin, and Fip1 (S. Chan et al., 2011; Schönemann et al., 2014). Of these proteins, two (CSPF30 and WDR33) recognize and bind the AAUAAA poly(A) signal and, through other members of the complex, initiate cleavage (S. L. Chan et al., 2014; Clerici et al., 2018; Sun et al., 2018). Although not as clearly defined as in metazoans, A - rich motifs in budding yeast such as AAGAA play an analogous role as poly(A) signals (Gross & Moore, 2001; Hill et al., 2019; Kumar et al., 2019).

In multiple species, the AAUAAA hexamer is insufficient to direct cleavage, and additional auxiliary sequences within the nascent transcript strengthen the poly(A) signal to promote accurate cleavage and polyadenylation (Birse, 1997; Sheets et al., 1990). In metazoans, there are two major auxiliary elements: upstream U-rich motifs and downstream U- and UG-rich motifs. The most highly enriched U-rich motif is a UGUA tetramer that is recognized by proteins in the <u>C</u>leavage <u>f</u>actor <u>Im</u> (CFIm) family (Brown & Gilmartin, 2003; Venkataraman, 2005). U and GU-rich sequences downstream of the cleavage site are recognized by <u>C</u>leavage <u>s</u>timulation factor proteins (CstF) that also help to strengthen the poly(A) signal and direct the endonuclease CPSF73 for cleavage of the nascent RNA (Hu et al., 2005; Mandel et al., 2006; Sullivan et al., 2009; Takagaki & Manley, 1997). In yeast, a similar set of auxiliary elements also help define cleavage sites (Baejen et al., 2014; Dichtl, 2002).

Despite our deep knowledge of cleavage and polyadenylation in metazoans and yeast, little is known about the sequences and complexes involved in this process for other eukaryotes. For example, there are over 200,000 species of protists, but only for kinetoplastids such as *Trypanosoma* and *Leishmania* do we have a clear image of how pre-mRNAs are processed (Clayton & Michaeli, 2011; X.-Q. Li & Du, 2014). Kinetoplastids are unique among eukaryotes in that they transcribe genes as polycistronic mRNAs that are cleaved post-transcriptionally to generate individual transcripts (Campbell et al., 2003; Clayton, 2019). Although *Trypanosomes* contain most of the conserved eukaryotic cleavage and polyadenylation proteins, the site of cleavage is established by the trans-splicing of the upstream gene and is not dependent on a specific motif (Clayton, 2013, 2019; Hendriks et al., 2003; Koch et al., 2016). This unusual mechanism speaks to the diversity of pre-mRNA processing in eukaryotes and highlights the gap in knowledge for the mechanisms controlling cleavage and polyadenylation in protists. As such, our knowledge of the conservation of the sequences, proteins, and mechanisms involved in cleavage and polyadenylation is limited, as is our knowledge of how this process has evolved and diverged.

One protist that has attracted interest is *Giardia lamblia*. A human parasite, *G. lamblia* is the causative agent of giardiasis, one of the most common intestinal diseases world-wide (Ankarklev et al., 2010). The *Giardia* clade encompasses multiple species that colonize the intestines of a variety of animals. It is also deeply branching, diverging from other eukaryotes over a billion years ago. From the perspective of gene regulation, *G. lamblia* has several deviations from model organisms that make it an ideal candidate to understand the evolution and diversification of fundamental machinery. First, previous work has suggested that the *3′* UTRs of *G. lamblia* are unusually short, with a median of less than 100 nucleotides (Franzén et al., 2013). This observation has raised basic questions about the potential for 3′ UTR-mediated regulation of RNA decay and translation in this organism. Second, consistent with short UTR regions, the genome of *G. lamblia* is generally very compact with only eight genes that contain introns and five that undergo trans-splicing, while the number of protein-coding genes is between 5000 to 9000, depending on genome annotation (Xu, Jex, et al., 2020). Third, *G. lamblia* has streamlined machinery for transcription (Best, 2004; Morrison et al., 2007), splicing (Iyer et al., 2019; Nixon et al., 2002), and translation (Eiler et al., 2020; L. Li & Wang, 2004), and lacks many protein components that are essential for viability in most other eukaryotes, such as the translation initiation factor eIF4G (L. Li & Wang, 2004). Finally, *G. lamblia* exists in two forms — as a dormant and hardy cyst and as an infectious trophozoite—making it a potential model system to investigate how cell state and developmental transitions affect gene expression. However, despite growing interest in *G. lamblia*, fundamental aspects of pre-mRNA processing, including the identity of its poly(A) signal and auxiliary elements, remain unknown.

To provide an initial genome-wide characterization of *G. lamblia* 3′ end processing, we generated high quality *G. lamblia* 3′ UTR annotations using two orthogonal high-throughput sequencing methods. Using these data, we identified the *G. lamblia* poly(A) signal: AGURAA (where R indicates a purine). Unlike yeast, *G. lamblia* uses a specific hexamer as its poly(A) signal, but this sequence differs from that of metazoans at the second position, using a G rather than an A. This unusual poly(A) signal has shaped the *G. lamblia* genome, with the hexamer depleted in coding regions while, at times, overlapping with stop codons. We looked for known auxiliary sequences surrounding the cleavage sites, but found little evidence of these elements playing an important role. Many of the proteins that would recognize auxiliary sequences are also absent, and together our results suggest that *G. lamblia* has pared-down pre-mRNA processing machinery or that the sequences and complexes have diverged to the point where they are difficult to identify. Finally, we identified 133 genes with more than one cleavage site. These results substantially increase the number of alternative polyadenylation events in *G. lamblia* (previously two (Mok et al., 2005; Que et al., 1996)). Our results therefore suggest that, despite simplified cleavage and polyadenylation machinery, 3′ end formation is an important, and as-yet underappreciated, mechanism for regulating gene expression in *G. lamblia*.

## RESULTS

### Characterization of *G. lamblia* mRNA 3′ ends at nucleotide resolution

To directly annotate *G. lamblia* 3′ UTRs, we began with a commercially available 3′-end sequencing method called QuantSeq, which maps 3′ ends of polyadenylated RNA with nucleotide resolution. We generated two replicate libraries using trophozoite RNA. Putative cleavage sites were defined by identifying the positions of read peaks downstream of annotated coding regions. Peaks that were within 10 nucleotides of each other were merged into a single site. We then filtered the sites to only include those where the predicted 3′ UTR showed at least 90% overlap between biological replicates. In some cases, these libraries led to multiple putative cleavage sites per gene, some of which were found tens of thousands of nucleotides away from the nearest open reading frame.

Given the known artefacts of this method (such as internal priming (Adiconis et al., 2013; Nam et al., 2002)) and the challenge of working with a relatively poorly annotated genome, we next used an orthogonal method to validate cleavage sites predicted by the 3′-end seq libraries. We directly sequenced *G. lamblia* RNA in duplicate using Oxford nanopore technology (ONT) and obtained 1.1 million total reads. ONT sequencing yields long reads (average length in our libraries: 940 bp) that enabled us to unambiguously develop precise gene models and thus enhance the transcriptomic map of poorly annotated species like *G. lamblia*.

We used two criteria for ONT read inclusion: reads were required to (1) have a poly(A) tail of at least 30 nucleotides (suggesting that they were derived from mature transcripts, Figure S1A, B) and (2) extend into any part of the open reading frame of the nearest gene (suggesting that they were genuine transcripts from that gene). To validate a cleavage site, we required that it was included in our QuantSeq dataset and had at least one read in either of the two replicate ONT libraries. This method allowed us to remove cleavage sites resulting from internal priming as well as misassigned sites, such as those that belonged to previously unannotated genes (Figure 1A). With this combined approach, we were able to identify 2764 cleavage sites across 2630 genes (which we will refer to as “validated cleavage sites”, Figure S1C).

**Figure 1.**
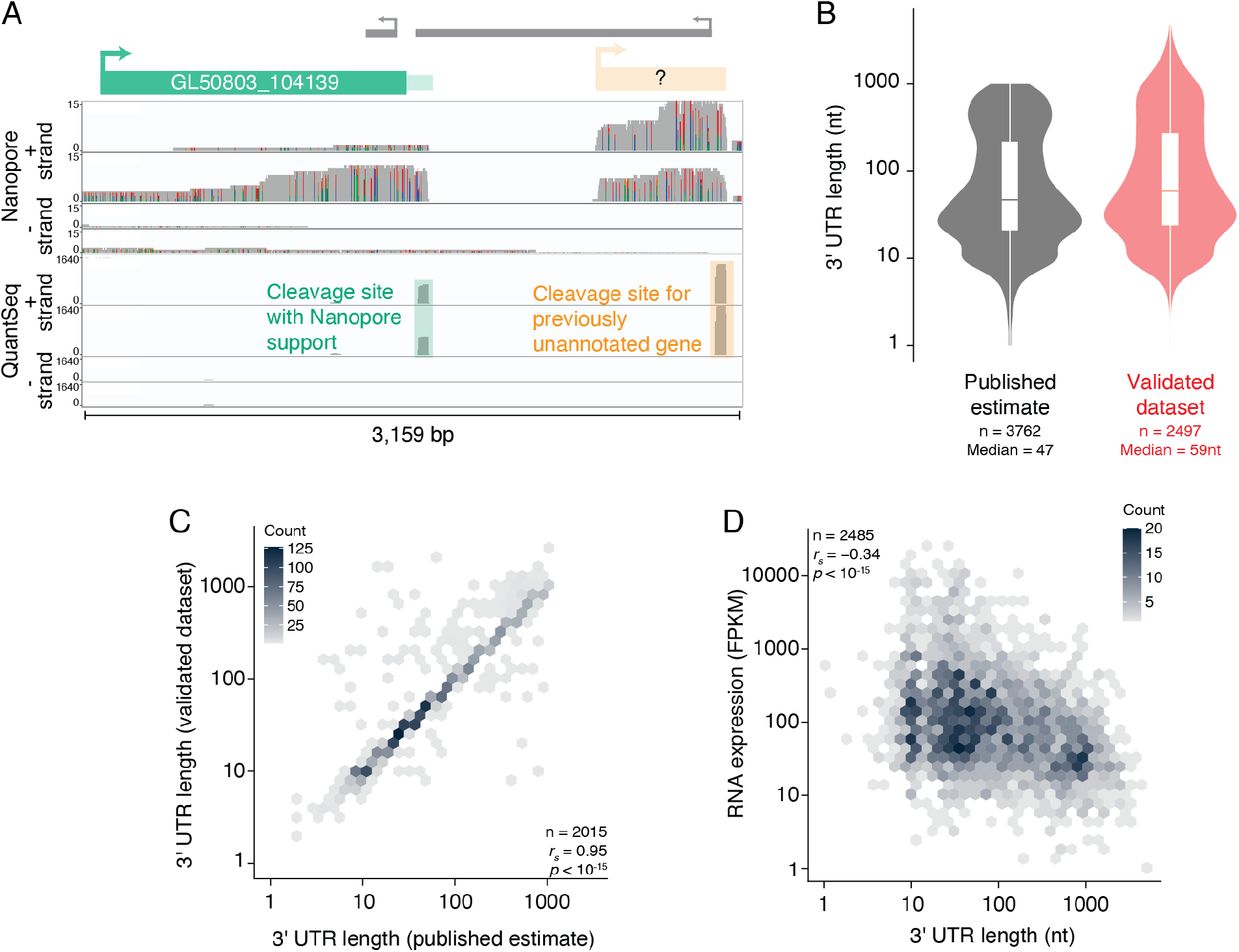
Characterization of G. lamblia 3′ ends at nucleotide resolution. (A) Genome browser image looking at the 3′ end of GL50803_104139 and displaying coverage of ONT libraries (top) and 3′-end libraries (bottom). Of the two cleavage sites predicted by the 3′-end libraries, one is supported by the ONT libraries (green box), while the other appears to belong to a previously unannotated transcript (orange box). (B) Distribution of 3′ UTR lengths in previously published work ((Franzen et al., 2013), left) and this study (right). (C) Hexagonal heatmap comparing previously published estimates of 3′ UTR lengths (x-axis) and the new dataset from this study (y-axis). (D) 3′ UTR length is negatively correlated with expression. Shown is a hexagonal heatmap comparing 3′ UTR length (this study) and mRNA expression in Fragments Per Kilobase of transcript per Million mapped reads (FPKM, from accession number GSE158187).

To further validate our results, we first compared them to the 3′ UTR length of genes that had previously been determined experimentally. For instance, cyst wall protein 1 (CWP1) has been described as having a 36-nucleotide 3′ UTR, and, consistent with these reports, our measurement gave 37 nucleotides (Hehl et al., 2000). Likewise, we found that NADP-specific glutamate dehydrogenase (GDH) has a 22-nucleotide long 3′ UTR (Figure S1D), consistent with previous predictions (Yee & Dennis, 1992). Thus, by using a combination of 3′-end seq and long-read sequencing, we generated a high-confidence dataset of validated cleavage sites for thousands of *G. lamblia* genes.

We also compared our annotations with those previously predicted on a genome-wide scale (Franzén et al., 2013). The 3′ UTR lengths generated by our approach had a median of 59 nucleotides and a similar distribution to previous predictions (Figure 1B). Although these previous estimates and our own annotations were highly correlated (Spearman *r [r_s_]* = 0.95, *p* < 10^-15^, Figure 1C), for 693 genes our experimentally determined 3′ UTRs were longer than the previous predictions, highlighting the power of our approach. We also observed a significant negative correlation between 3′ UTR length and mRNA expression (Figure 1D, Spearman *r [r_s_]* = −0.34, *p* <10^-15^), as has been observed in other organisms (Mayr, 2017). This result raises the possibility that 3′ UTRs, despite their short length, may carry sufficient regulatory potential to modulate mRNA stability, although the associated mechanisms are unknown.

### *G. lamblia* uses an unusual poly(A) signal

From our list of validated cleavage sites, we next asked which poly(A) signals, if any, *G. lamblia* uses. As a first approach, we looked at the frequency of each nucleotide in a 60-nucleotide window centered on the validated cleavage sites (Figure 2A). We noticed A-richness approximately 10 nucleotides upstream of the cleavage site, as well as a distinct A-peak directly downstream. There was also an enrichment of U nucleotides both up and downstream of this region, similar to that seen in other organisms ((Tian et al., 2005; Tian & Graber, 2012), Figure S2A). These results suggest that *G. lamblia* has sequence preferences for defining cleavage sites.

**Figure 2.**
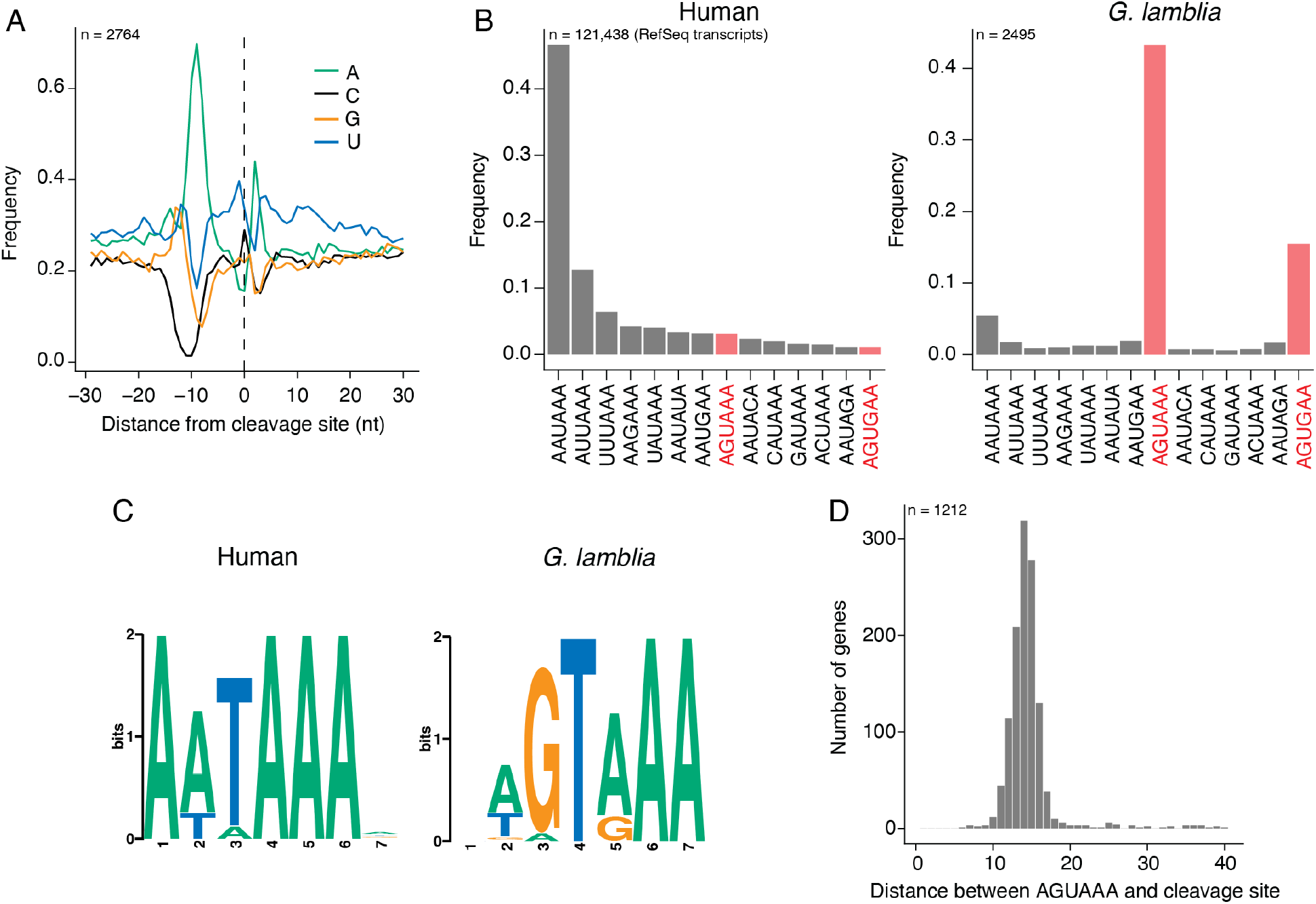
*G. lamblia* uses an unusual poly(A) signal. (A) Nucleotide frequency in the 60-nucleotide window centered on all 2860 validated cleavage sites from this study. (B) Frequency of common poly(A) signals identified in studies of human transcripts (Beaudoing, 2000). Sequences 30 nucleotides upstream of cleavage sites from the human RefSeq annotations and validated *G. lamblia* sites from this study were used to search for common motifs. Plotted is the frequency of each signal in human (left) and *G. lamblia* (right). (C) MEME analysis of upstream sequences. The same sequences as in (B) were uploaded to the meme-suite and a search was conducted for enriched hexamers. Shown is the top motif for human (left) and *G. lamblia* (right). (D) For all validated cleavage sites containing an AGUAAA motif in the last 40 nucleotides of the mRNA, this bar graph shows the distance between the motif and the end of the read. Distances are counted from the first A of the motif.

We next wanted to define the precise poly(A) signal used in *G. lamblia*. To do so, we focused on genes with only one validated cleavage site and counted the occurrences of hexameric motifs previously identified in humans (Beaudoing, 2000). When we performed this analysis on human RefSeq transcript annotations, as expected AAUAAA is the most abundant polyadenylation signal (Figure 2B). Surprisingly, distinct, but related, motifs were the most highly enriched in our *G. lamblia* dataset: AGUAAA and AGUGAA were found in 45% and 15% of genes, respectively. In contrast, AAUAAA was used only rarely, occurring in 5% of genes.

As an independent approach, we also searched for 6 nucleotide motifs occurring within the first 30 nucleotides upstream of human and *G. lamblia* cleavage sites using the MEME package (Bailey et al., 2009). This unbiased approach confirmed the strong enrichment for the G nucleotide at position 2 of the poly(A) signal and the strong preference for a purine at position 4 (Figure 2C). Our identified poly(A) signal is also consistent with early studies of individual *G. lamblia* transcripts that suggested an AGURAA motif as the polyadenylation signal (Peattie et al., 1989; Que et al., 1996; Yee & Dennis, 1992) — an observation we have now confirmed on a genomewide scale.

Interestingly, although metazoan poly(A) signals are usually found 10 to 30 nucleotides upstream of the cleavage site (Kumar et al., 2019; Figure S2B), *G. lamblia* signals tended to be closer to the cleavage site (Figure 2D). In over 90% of genes with an AGUAAA signal, the motif was less than 20 nucleotides from the cleavage site, and the most common distance was 13–15 nucleotides, an observation consistent with the general compactness of the *G. lamblia* genome.

### Implications of unusual poly(A) signal on the *G. lamblia* genome

We next wished to investigate how the unusual poly(A) signal has shaped the *G. lamblia* genome. First, given that AGUAAA and AGUGAA are poly(A) signals, we would expect them to be depleted in open reading frames as their presence early in the transcript would lead to premature cleavage. To test this prediction, we counted the occurrence of both motifs and compared them to the frequency of their shuffled sequences (*e.g*. AAUAGA). We found that AGUAAA is strongly depleted in open reading frames compared to the shuffled sequences while the depletion of AGUGAA was more modest, consistent with the prediction that AGUAAA is the preferred signal (Figure 3A-B).

**Figure 3.**
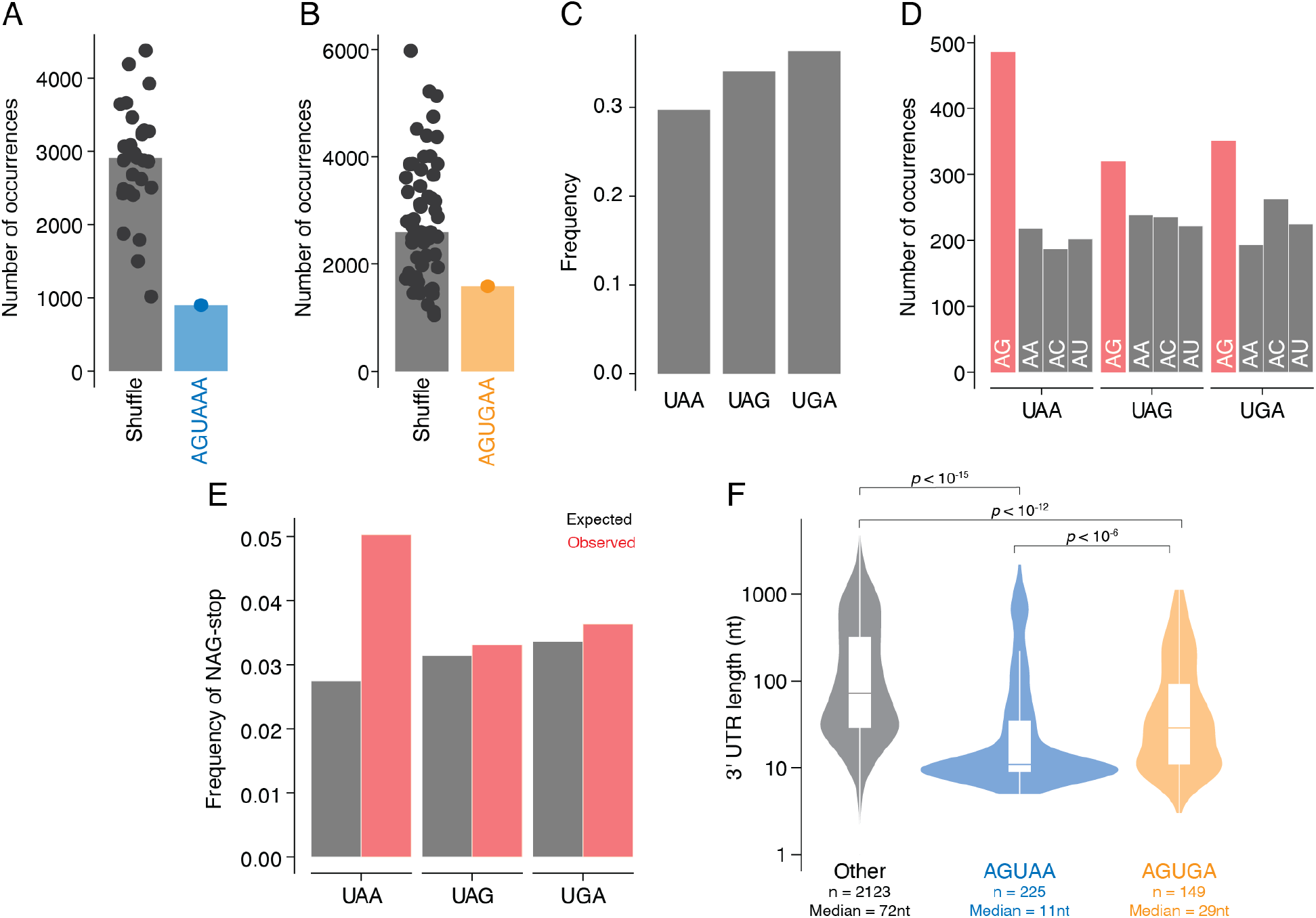
Implications of unusual poly(A) signal on *G. lamblia* open reading frames. A) Open reading frames are depleted for *G. lamblia’s* poly(A) signal. Open reading frame sequences were used to count the occurrence of AGUAAA vs all shuffled versions of the motif. (B) As in A, but with the AGUGAA poly(A) signal. (C) Frequency of stop codons across all annotated *G. lamblia* open reading frames. (D) Nucleotides preceding a stop are enriched for AG over other AN dinucleotides. For each stop codon, this bar graph shows how many were preceded by the different AN dinucleotide sequences. (E) As in D, but comparing expected vs observed frequencies. The expected frequency for each sequence context was calculated from the total frequency of each codon across all open reading frames. (F) Distribution of 3′ UTR lengths for genes where there is no overlap of poly(A) signal and stop codon (left), genes where there is an AG dinucleotide preceding a UAA stop codon (middle), and genes where there is an AG preceding a UGA stop codon (right).

We then investigated the relationship between poly(A) signals and stop codons following a recent study of *Giardia muris* that reported that many genes have an overlap between these signals (Xu, Jiménez-González, et al., 2020). Genomic analysis of *Spironucleus salmonicida*, another diplomonad, also suggests a strong “dual use” of poly(A) signals as stop codons (Xu et al., 2014). In the case of *S. salmonicida*, there is the stop codon (UGA) is predominantly used throughout the genome, overlapping with a predicted an AG<u>UGA</u> poly(A) signal (Xu et al., 2014). Given the short length of 3′ UTRs in *G. lamblia*, we wondered whether this overlap of signals might also occur here. We first calculated the frequency of each stop codon across all open reading frames. We did not observe a strong preference for any sequence, and UAA (which would allow for an AG-UAA motif) was the least abundant of the 3 stop codons (Figure 3C). We next looked more closely at the nucleotides preceding the stop codon and asked whether there was a preference for AA, AU, AC or AG. Of these, only a (N)AG sequence in front of the stop codon will allow for a dual AG<u>UAA</u>A or AG<u>UGA</u>A poly(A) signal/stop codon combination. Although there was no enrichment for the UAA stop codon itself in open reading frames, it was much more likely to be preceded by an NAG codon than the other codons. We also observed a modest preference for AG dinucleotides preceding the other two stop codons (Figure 3D). In contrast, AA dinucleotides showed no such preference, providing an additional line of support that *G. lamblia* does not use the AAUAAA hexamer.

Two alternative models could explain the nucleotide bias in the codon preceding the stop codon: the first is that NAG-UAA and NAG-UGA represent genuine poly(A) signals and the second is that their presence is simply a consequence of codon usage or amino acid preferences. To distinguish between these possibilities, we compared the expected and observed frequencies of NAG sequences preceding the stop codon. Consistent with NAG-UAA serving as a dual poly(A) signal/stop codon, this dicodon occurred more frequently than expected based on the frequencies of either alone. The same was not true for NAG-UGA (Figure 3E). To investigate this further, we looked at the 3′ UTR lengths of genes with the potential dual use AG-UAA or AG-UGA stop codons. Compared with other genes, 3′ UTR lengths were shorter for both AGUAA- and AGUGA-ending transcripts (*p*<10^-15^ and *p*<10^-12^, respectively; Figure 3F). In the case of AGUAA, the median length was 11 nucleotides, which is in the window of distances between genuine poly(A) signals and cleavage sites. These analyses indicate that NAG-URA sequences can act as genuine dual-use stop codons and poly(A) signals. In other words, in *G. lamblia*, stop codons have acquired the ability to also act as poly(A) signals for ~15% of genes. However, this dual usage has not reached the levels predicted in *G. muris* and *S. salmonicida*, suggesting that this aspect of genome organization is evolving relatively rapidly within the diplomonad order.

### Eukaryotic auxiliary elements are poorly enriched around *G. lamblia* cleavage sites

We have an advanced understanding of the sequences and proteins involved in the recognition of polyadenylation signals and auxiliary elements in other eukaryotes. In metazoans, there are three main complexes that recognize the polyadenylation signal, upstream U-rich motifs and downstream U- and UG-rich motifs: CPSF, CFIm and CstF complexes, respectively (Brown & Gilmartin, 2003; Kumar et al., 2019; Takagaki & Manley, 1997). However, it is completely unknown whether *G. lamblia* also makes use of auxiliary elements to define cleavage sites.

To investigate whether these sequences were conserved in *G. lamblia*, we began by searching for orthologs to the associated proteins. Although we readily identified candidates for the CPSF complex (which recognizes the poly(A) signal), we found only low-confidence candidates for members of the CstF complex (which recognizes downstream U-rich motifs), and we were unable to identify orthologs for the CFlm proteins (which recognize upstream U-rich motifs and UGUA; Figure 4A, Supplemental Table 2).

**Figure 4.**
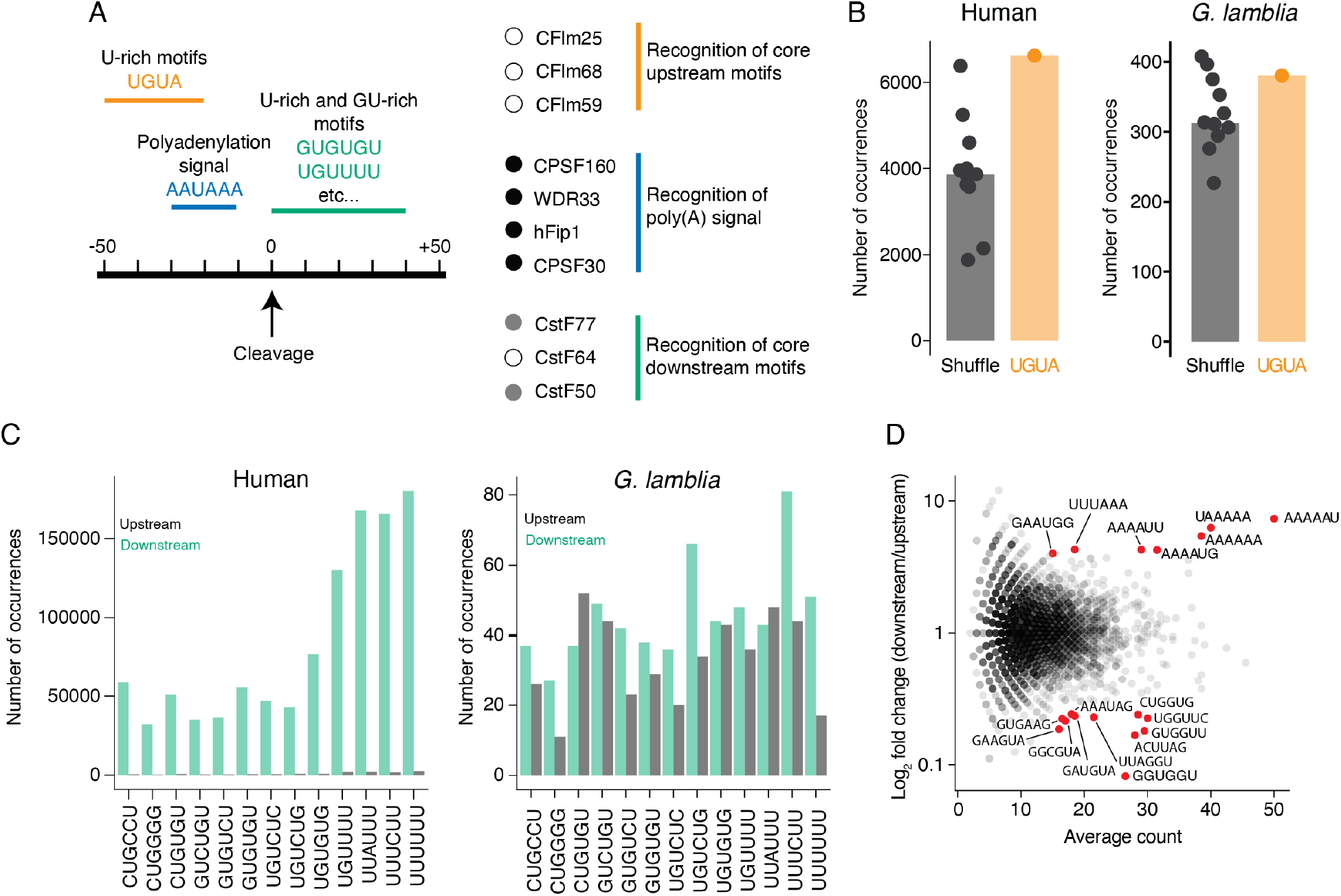
Conserved auxiliary elements are poorly enriched around *G. lamblia* cleavage sites. (A) Conserved pre-mRNA processing proteins and the sequences they recognize. The left panel shows the location and motifs of key sequences found around human cleavage sites. Right panel shows the human orthologs of core processing complexes for the recognition of poly(A) signals and surrounding sequences. Dots indicate whether an ortholog was readily identifiable in *G. lamblia* (black circle), whether ortholog identification was ambiguous (grey circle), or whether no orthologs were found (white circle). (B) The conserved UGUA motif is not enriched upstream of *G. lamblia* cleavage sites. Sequences 20 to 50 nucleotides upstream of cleavage sites were used to count the frequency of UGUA or shuffled versions of the motif. Plotted is the number of times each motif was found in human (left) and *G. lamblia* (right) sequences. (C) GU-rich elements are not enriched downstream of *G. lamblia* cleavage sites. Sequences 40 nucleotides up and downstream of human and *G. lamblia* cleavage sites were used to count the occurrence of U- and GU-rich motifs enriched downstream of strong human cleavage sites (Hu et al., 2005). Plotted is the frequency of each motif upstream (grey) or downstream (green) of human (left) and *G. lamblia* (right) cleavage sites. (D) MA plot of enriched and depleted 6-mer sequences around polyadenylation signals. All single cleavage site genes from our dataset that contain an AGUAAA were selected for this analysis. Sequences 50 nucleotides upstream and downstream of the signal were used to search for all possible 6-nucleotide motifs. Plotted is the average count of each motif versus its enrichment in downstream sequences. Red dots are motifs that showed at least a 4-fold enrichment or depletion in downstream regions and with an average count of at least 15 occurrences.

We next looked at the sequences surrounding *G. lamblia* cleavage sites to investigate the extent to which the corresponding recognition sequences of these complexes were enriched. We first looked at the sequences 20 to 50 nucleotides upstream of the cleavage sites where the highly conserved UGUA motif is found in other eukaryotes (Brown & Gilmartin, 2003; Millevoi & Vagner, 2010). By counting the number of occurrences of UGUA as well as shuffled versions of the motif, we observed a strong preference for UGUA in the human genome, as expected. In contrast, we saw only a slight enrichment in *G. lamblia* (Figure 4B). Consistent with this result, when we performed an unbiased motif search using MEME, no sequences were enriched in this region (data not shown). This poor sequence conservation, combined with our inability to identify any CFlm orthologs, suggest that upstream motifs either do not play a role in the processing of *G. lamblia* transcripts or are sufficiently divergent as to preclude identification.

Next, we looked for downstream auxiliary elements. In other organisms, these downstream elements lack a consensus motif, but rather are generally U-rich. Thus, we looked for hexamers that were enriched around strong poly(A) sites in human sequences (Hu et al., 2005). As expected, we found that these sequences were highly enriched in regions downstream of cleavage sites in humans, but almost completely absent upstream. In contrast, in *G. lamblia* the sequences were equally present either side of cleavage sites (Figure 4C), which suggests that *G. lamblia* does not use conserved downstream auxiliary elements. However, because the strong U bias observed downstream of the cleavage site in metagene analyses (Figure 2A) and the ambiguous presence of putative CstF orthologs raise the possibility that instead divergent *cis*-elements and proteins may help define genuine cleavage sites, we turned to an unbiased approach to look for enriched motifs. For each gene containing a single cleavage site and an AGUAAA poly(A) signal, we searched for all possible 6-nucleotide motifs in the 50 nucleotides upstream and downstream of the signal. We found an enrichment for A-rich and AU-rich motifs in the downstream regions, and a depletion of more canonical GU-rich motifs (Figure 4D). These results support our observation that any sequences that may help strengthen poly(A) signals in *G. lamblia* have diverged substantially from those found in classical model eukaryotes.

### Evidence of alternative polyadenylation in *G. lamblia*

As mentioned above, when annotating cleavage sites, we unexpectedly found 133 genes showing evidence of alternative polyadenylation (Figure 5A). Although there are two previously described examples of alternative polyadenylation in the *G. lamblia* literature (Einarsson et al., 2016; Mok et al., 2005; Que et al., 1996), this phenomenon was not believed to be widespread. Our results indicate that alternative polyadenylation is much more common than current models suggest (Figure S3A).

**Figure 5.**
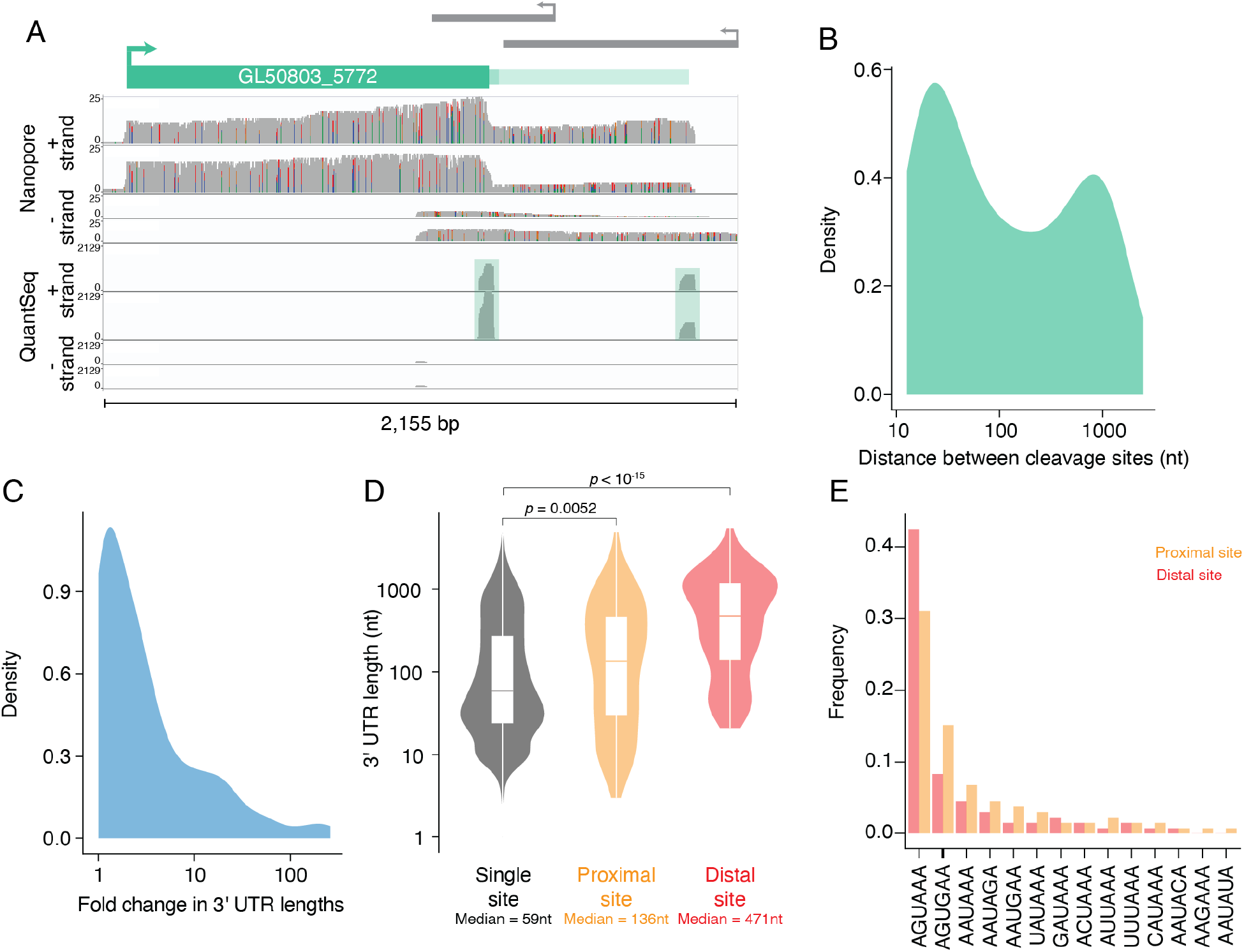
Evidence of alternative polyadenylation in *G. lamblia*. (A) Genome browser image looking at the 3′ end of GL50803_5772 and displaying cover-age of ONT libraries (top) and 3′-end libraries (bottom). Both methods support the presence of two distinct cleavage sites for the gene. (B) Density plot showing the distribution of lengths between proximal and distal cleavage sites for the genes that have more than one cleavage site. The median is 81 nucleotides. (C) Density plot showing the fold change in 3′ UTR length between distal and proximal cleavage sites. Median is a 2.18-fold change. (D) Distribution of 3′ UTR lengths for genes with a single cleavage site (left), the proximal sites for APA genes (middle), and the distal sites (right). (E) Poly(A) signal usage in APA genes. Sequences 30 nucleotides upstream of proximal and distal cleavage sites were used to search for the motifs described in Figure 2B. Plotted is the frequency of each motif across proximal (orange) and distal (red) cleavage sites.

The majority of these alternative cleavage sites were within 100 nucleotides of each other, although for 20 genes the distal cleavage site was over 1000 nucleotides downstream from the proximal one (Figure 5B). Nonetheless, given the short length of 3′ UTRs in *G. lamblia*, in 53% of cases, usage of the distal site more than doubled the amount of regulatory sequence (Figure 5C, Figure S3B). Interestingly, usage of the proximal site resulted in longer 3′ UTRs than in the rest of the transcriptome (Figure 5D: 135 nucleotides vs 59 nucleotides, *p* = 0.0052). In humans, proximal sites often use “weaker” poly(A) signals than distal sites (Hu et al., 2005; Legendre & Gautheret, 2003), and so we asked whether the same held true in *G. lamblia*. We found that distal cleavage sites are more likely to use AGUAAA and that proximal sites have a higher frequency of alternate signals such as AGUGAA (Figure 5E).

The presence of APA, as well as the generally longer 3′ UTR lengths observed, suggested that the regulation of this subset of genes may be biologically important. We observed a slight difference in overall expression between genes that had a single or multiple cleavage sites (median FPKM: 86 and 59.2, respectively; *p* = 0.00024; Figure S3C), although there was no difference in poly(A)-tail lengths (*p* = 0.39; Figure S3D). We also performed a gene ontology enrichment analysis, but no significant processes were enriched in genes undergoing alternative polyadenylation. We suspect that this result may be due to the fact that over 50% of genes are uncharacterized in *G. lamblia*, which limits the power of these approaches. Indeed, 12 of the alternative polyadenylation genes are described as ‘putative’, and 72 are hypothetical proteins or unspecified products. Nonetheless, two ribosomal protein genes (S4 and S28), as well as nine predicted kinases use alternative polyadenylation (Supplemental Table 1), raising the intriguing possibility that alternative polyadenylation may be important for the *G. lamblia* life cycle.

## DISCUSSION

Here, we empirically annotated the 3′ UTRs for 2630 expressed genes in *G. lamblia* using a combination of 3′-end short- and long-read sequencing. According to our RNASeq data, 6616 of the 9700 predicted coding genes in the genome annotation used for this study are expressed at an FPKM of 10 or higher. This indicates that we have annotated about 40% of the expressed transcriptome. Although one barrier to annotating the rest of the genome is low ONT sequencing depth (relative to short-read based sequencing) and the very low RNA expression of the remaining genes (average FPKM = 1.89), direct long-read RNA sequencing was nonetheless instrumental in overcoming some of the difficulties associated with the study of an organism whose genome remains relatively unannotated compared to traditional model systems. Critically, our use of ONT sequencing mitigated known issues with 3′ end short read sequences (Adiconis et al., 2013) and directly linked cleavage sites and open reading frames.

Our work confirms early hypotheses of an alternative poly(A) signal (Peattie et al., 1989; Que et al., 1996; Yee & Dennis, 1992), demonstrating that *G. lamblia* uses AGURAA on a genome-wide scale. Interestingly, the most frequent signal (AGUAAA) differs from the metazoan AAUAAA motif by only a single nucleotide, using a G at position 2 rather an A—but the two most common *G. lamblia* signals (AGURAA) are used only rarely in metazoans (Hu et al., 2005). One interesting question is how this divergent sequence is recognized. In metazoans, the poly(A) signal is recognized by CPSF30 and WDR33 (Casañal et al., 2017; S. L. Chan et al., 2014; Clerici et al., 2018). We were able to identify putative orthologs to these key players, but orthologs for supporting proteins such as CPSF-100 and Symplekin remain to be found (Supplemental Table 2). Identifying the appropriate orthologs and their sequence, structure, and biochemical preferences will be an important next step for understanding the basis of the unique *G. lamblia* poly(A) signal and its evolution.

Although starting with conserved eukaryotic sequences proved to be a good strategy when looking for polyadenylation signals, it was not the case for auxiliary elements. We were unable to find evidence of enrichment for any of the most common metazoan sequences that are found up or downstream of cleavage sites. It is therefore likely that any motifs outside the poly(A) signal used by *G. lamblia* to direct 3′-end processing have diverged significantly from those found in other eukaryotes, and their identification will likely require additional functional studies.

Finally, an unexpected finding from our study of 3′ UTRs is that 133 genes use alternative polyadenylation. Previous reports had identified only two cases (Mok et al., 2005; Que et al., 1996), suggesting that alternative polyadenylation was as rare as splicing in *G. lamblia*. Our results demonstrate that, contrary to this model, alternative polyadenylation is a more generally used mechanism, adding to the regulatory layers used by *G. lamblia*. Indeed, our results raise more intriguing questions about how cleavage and polyadenylation is regulated. For instance, how do these different 3′ UTR isoforms affect transcript stability and translation? Why do some genes use alternative polyadenylation, and not others? Previous reports have suggested that encystation impacts gene expression as well as cleavage and polyadenylation of individual genes (Einarsson et al., 2016; Mok et al., 2005; Que et al., 1996). An intriguing possibility is that alternative polyadenylation may be especially important during this process, or in the cyst itself (which is transcriptionally silent), and it will be exciting to explore this and other questions in the future.

## MATERIALS AND METHODS

### Trophozoite culture and RNA extraction

*G. lamblia* trophozoites (assemblage A, strain WB clone C6) were grown in modified TYS-33 media as per standard protocols (Keister, 1983). Cells were harvested by placing culture tubes on ice for 10 minutes, then spun down for 5 minutes at 800 x g at 4°C. Cell pellets were washed twice in 1xPBS. RNA was extracted from trophozoite pellets with hot acid phenol as previously described (Collart & Oliviero, 1993).

### RNA sequencing and analysis

Previously generated RNASeq libraries used in this study are available from the GEO (GSE158187). 3′-end libraries were generated with the QuantSeq 3′ mRNA-Seq Library REV kit from Lexogen (catalog #016) according to manufacturer’s protocol. Libraries were sequenced at the Genomics and Microarray Shared Resource at the University of Colorado Denver Cancer Center. All sequencing data generated in this study is available from the GEO, accession number GSE168675.

Nanopore libraries were prepared according to the Direct RNA sequencing protocol from ONT (SQK-RNA002). Because the lengths of poly(A) tails were unknown when we initiated this study, total RNA was used in place of poly(A) selected RNA. Libraries were sequenced on a FLO-MIN106 flow cell and minION sequencing device. Base calling was completed by the MinKNOW software (Nanopore) on default settings.

Adaptors were trimmed from 3′-end reads using Cutadapt v2.3. RNASeq and QuantSeq libraries were aligned using STAR 2.5.2a (Dobin et al., 2013). Nanopore libraries were aligned with minimap2 version 2.17-r974-dirty (H. Li, 2018). All libraries were mapped to the *Giardia lamblia* WBC6 genome version 50 downloaded from the GiardiaDB website on February 8, 2021 (https://giardiadb.org). Poly(A) tail lengths from the Nanopore libraries were measured using Nanopolish version 0.11.1 (Loman et al., 2015). Mapped nanopore reads were assigned to their corresponding gene using featureCounts version 2.0.0 (Liao et al., 2014).

### Identification and validation of cleavage sites

3′ UTRs were annotated by first identifying poly(A) sites. Poly(A) sites were mapped by identifying peaks of poly(A) reads that aligned downstream of coding regions but did not overlap the following gene. Potential poly(A) sites were filtered to only include those that have at least 10 reads. Sites that were within 10 nucleotides of each other were merged into a single peak.

For each putative cleavage site, a list of coordinates was generated that went 10 nucleotides up and downstream from the site. For each gene, ONT reads with 3′ ends that ended within the corresponding window were selected. Reads were then further filtered to keep only those that contained a poly(A) tail of at least 30 nucleotides and for which the 5′ end of the read fell within the open reading frame of the associated gene. Sites with at least one read from either replicate of the ONT libraries that satisfied all conditions were kept as validated sites. Analyses and plotting were performed in R version 4.0.3 and Python version 3.8.3 from in-house scripts. All genome browser images were generated with IGV version 2.8.10.

### Unbiased motif analysis

Motif-based sequence analysis was done using the MEME suite software at meme-suite.org (Bailey et al., 2009). We searched for a maximum of 3 motifs on the given strand only with minimum and maximum motif lengths of 6 and 50 nucleotides, respectively.

### Ortholog identification

Human protein sequences were used to search for orthologs in *G. lamblia* by BLAST search. Where it was difficult to identify the most likely ortholog among the search results, the yeast protein sequence was used for a complimentary search. Searches were conducted on giardiadb.org. For CPSF160 and WDR33, human proteins containing similar domains were used to perform a multiple sequence alignment, which was then used to generate a Hidden Markov Model. We then initiated a search across the *G. lamblia* proteome in search of proteins that have a similar domain and sequence.

## ACKNOWLEDGEMENTS

We thank Dr. Lori Passmore and Vytaute Boreikaite for insightful discussions about the CPSF complex. We thank members of the Rissland, Jagannathan, Bentley, Mukherjee, and Taliaferro labs for helpful and thoughtful discussions during our weekly lab meetings. We thank Dr. Staffan Svärd for assistance with *G. lamblia*. We are grateful to Dr. Passmore, Dr. Bentley, Dr. Mukherjee and Dr. Ramachandran for their feedback on this manuscript. This work was supported by NIH grants R35GM128680 (OSR) and the RNA Bioscience Initiative. ARJ is supported by the Australian National Health and Medical Research Council L1 Investigator grant (APP1194330) and the Walter and Eliza Hall Institute of Medical Research receives support through the Victorian State Government Operational Infrastructure Support and Australian Government National Health and Medical Research Council Independent Research Institute Infrastructure Support Scheme.

## SUPPLEMENTAL TABLE LEGENDS

**Supplemental Table 1. Validated 3′ UTR sites in *G. lamblia*.** Table of all validated 3′ UTR lengths generated in this study. Genes are in separate sheets according to the number of cleavage sites.

**Supplemental Table 2. Predicted orthologs for cleavage and polyadenylation proteins in *G. lamblia*.** List of human cleavage and polyadenylation factors and putative *G. lamblia* orthologs. Human proteins were taken from (Tian and Graber, 2012). Orthologs for *G. lamblia* were identified as described in methods.

---

**Supplemental figure 1.**
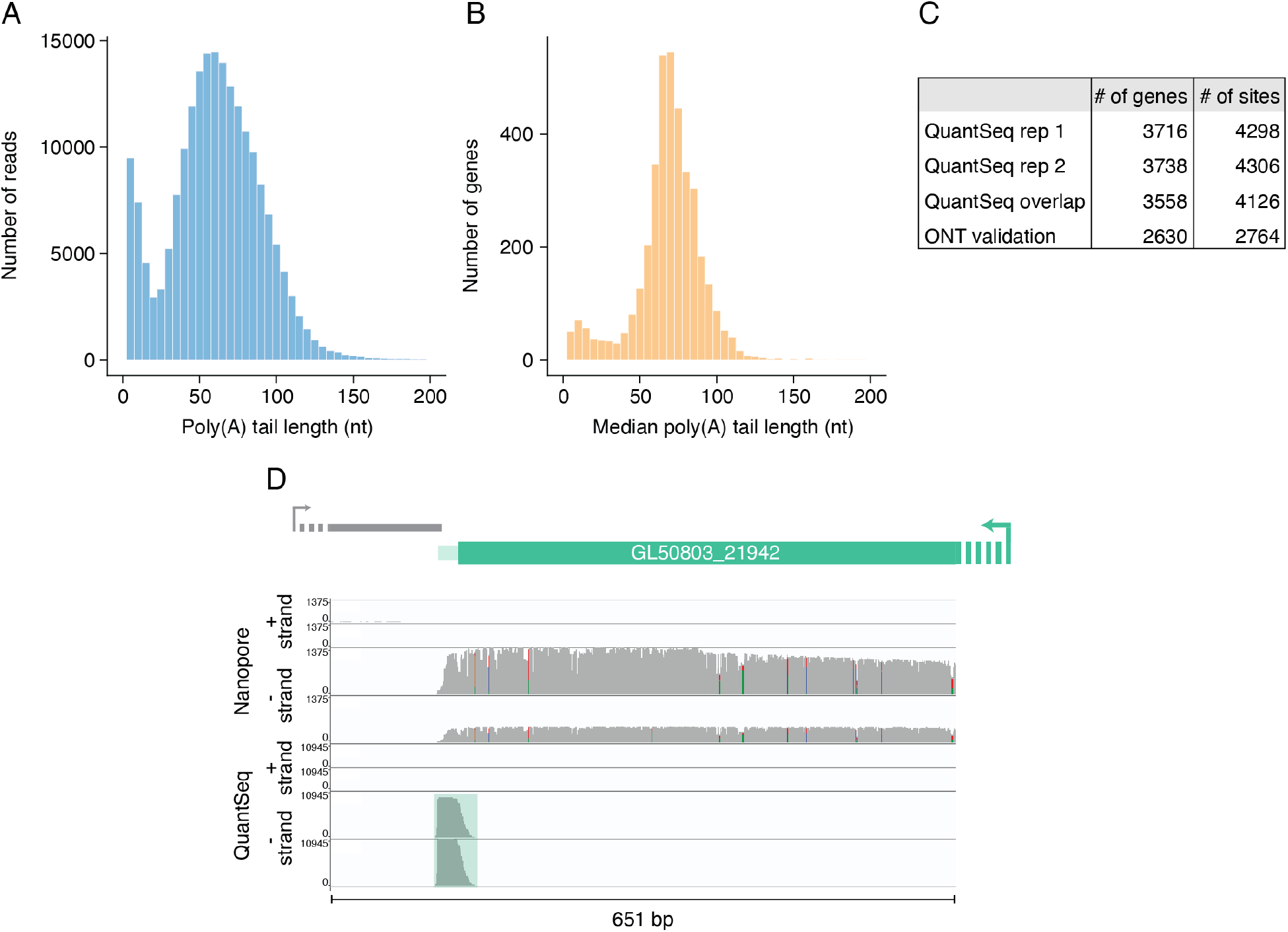
Poly(A) tail measurements allow for validation of cleavage sites. (A) Distribution of poly(A) tail lengths in ONT libraries. Histogram shows the poly(A) tail length of all reads that were assigned to a gene and for which a poly(A) tail was measured. Median is 60.5 nucleotides. (B) Reads in A were grouped by gene and plotted is the median poly(A) tail length for each gene. Ribosomal RNAs and tRNAs were removed for this analysis. The median length is 69.9 nucleotides. (C) Table showing the number of genes and sites for each step in our pipeline. “QuantSeq overlap” represents the 3′ UTRs that show at least 90% overlap between biological replicates. (D) Genome browser image looking at the 3′ end of GL50803_2I942 (NADP-specific glutamate dehydrogenase). 3′-end libraries and ONT libraries both show a cleavage site 22 nucleotides downstream from the stop codon, which is consistent with previous observations.

**Supplemental figure 2.**
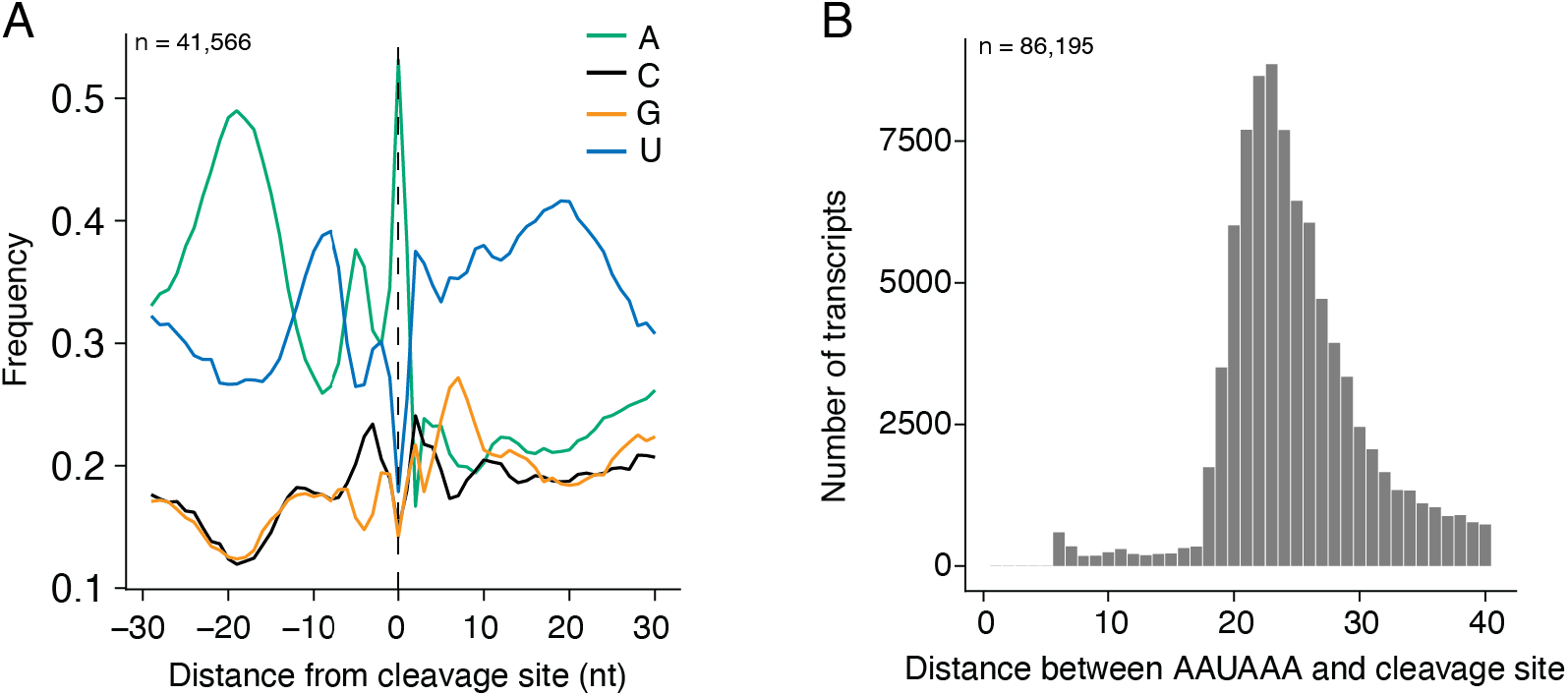
Cleavage and polyadenylation contexts in humans. (A) Nucleotide frequency surrounding human cleavage sites. Sequences surrounding the end coordinates of all rmRNAs in the RefSeq annotation were used to calculate nucleotide frequencies 30 nucleotides upstream and downstream of cleavage sites. (B) Distances between AAUAAA poly(A) signal and cleavage sites in human transcripts. mRNA sequences were downloaded from RefSeq annotations. All sequences with an AAUAAA in the last 40 nucleotides of the transcript were selected and the distance was calculated between the motif and the end of the mRNA. Distances are counted from the first A of the motif.

**Supplemental figure 3.**
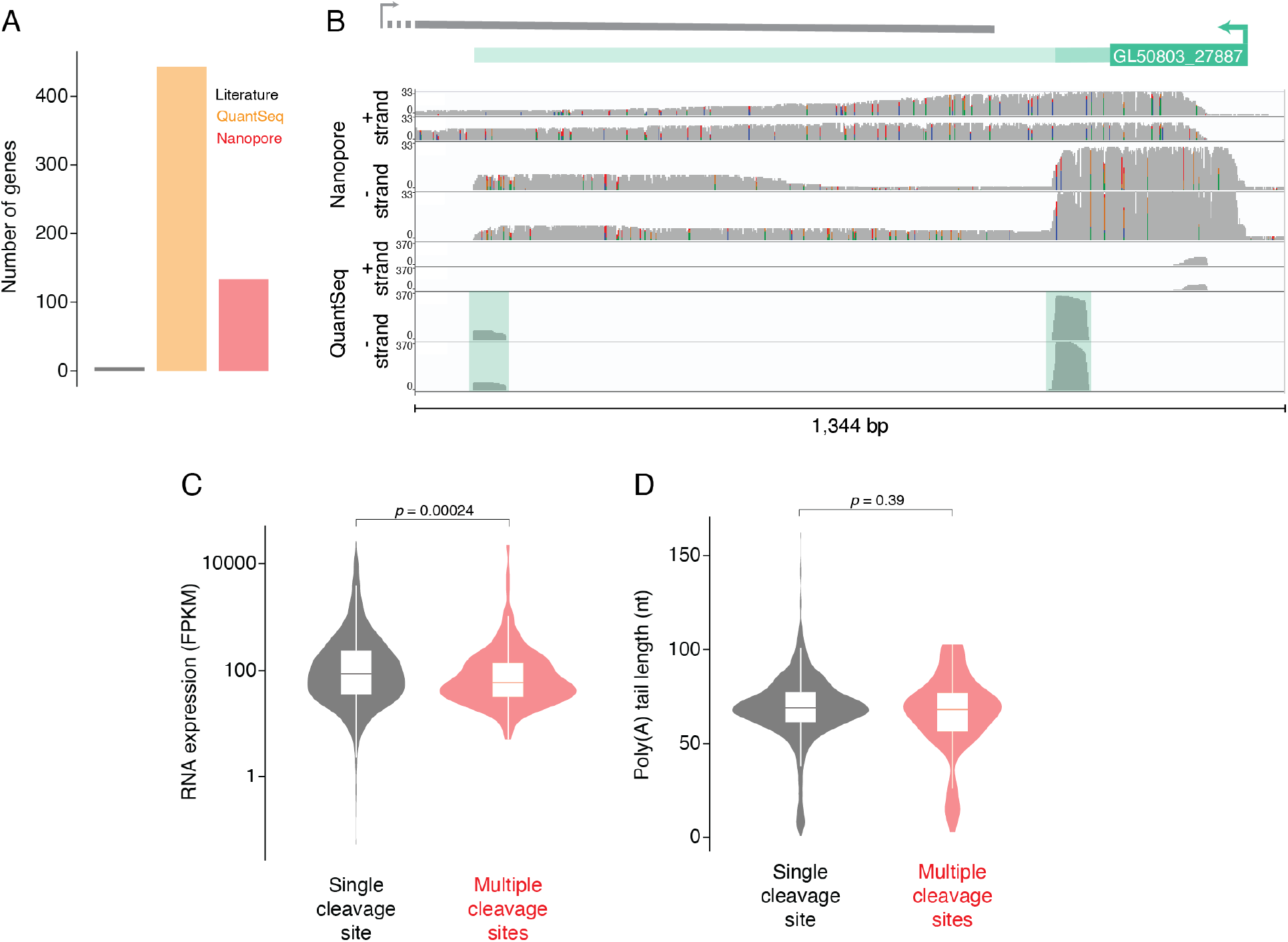
Gene differences between proximal and distal cleavage site usage. (A) The number of alternative polyadenylation events in *G. lamblia* is greater than previously thought. Bar plot shows the number of genes that were believed to contain more than one cleavage site based on previous literature, our 3′-end libraries alone (QuantSeq), and our validated list of cleavage sites (Nanopore). (B) Genome browser image showing a gene (GL50803_27887) with a short and long 3′ UTR measuring 86 and 984 nucleotides, respectively. (C) FPKM expression for genes with a single cleavage site (left) or multiple cleavage sites (right). Median values are 86 and 59.2, respectively. (D) Distribution of poly(A) tail lengths for genes with a single cleavage site (left) or multiple cleavage sites (right). Median values are 69 and 68.2 nucleotides, respectively.

